# Integrated protein and transcriptome high-throughput spatial profiling

**DOI:** 10.1101/2022.03.15.484516

**Authors:** Nir Ben-Chetrit, Xiang Niu, Ariel D. Swett, Jesus Sotelo, Maria S. Jiao, Patrick Roelli, Marlon Stoeckius, Dan A. Landau

## Abstract

Spatial transcriptomics and proteomics have independently transformed our understanding of complex biological processes; however, integration of these modalities is currently limited. To overcome this challenge, we developed **<u>S</u>**patial **<u>P</u>**r**<u>O</u>**tein and **<u>T</u>**ranscriptome **<u>S</u>**equencing (SPOTS) for high-throughput integration of transcriptome and protein profiling within the spatial context. Applying SPOTS to spleen and breast cancer samples revealed that spatially-resolved multi-omic integration provides a comprehensive perspective on key biological processes in health and disease.

## Main

The relationship between cells in their native microenvironment is critical to our understanding of mechanisms regulating diverse aspects of biology in health and disease. Single-cell genomics approaches (e.g., single-cell RNA-seq) have substantially increased the throughput and granularity of our ability to study biological processes in tissues^1,2^. Nonetheless, the spatial context is lost during tissue dissociation required for single-cell experiments. In contrast, spatial transcriptomics (ST) enables in-depth molecular characterization of tissues in a spatially-informed manner^3^. However, many cutting-edge methods, including High-Definition Spatial Transcriptomics (HDST)^4^, Slide-seqV2^5^, and Seq-Scope^6^, are currently limited to unimodal transcriptomic characterization. On the other hand, DBiT-seq combines protein detection with whole transcriptome measurement to further increase granularity; however, it relies on a sophisticated microfluidics apparatus that limits its widespread use^7^. Other recent advancements have incorporated extracellular protein detection through measurement with TotalSeq™ A antibodies into the broadly available Visium protocol^8,9^; however, they either rely on manual microarray design and automated robotic systems that are not readily accessible^8^, showed limited sensitivity for detecting the expected variability of proteins of interest^9^, and have not been demonstrated on highly fibrotic tissue such as solid tumors.

To address these challenges, we developed **<u>S</u>**patial **<u>P</u>**r**<u>O</u>**tein and **<u>T</u>**ranscriptome **<u>S</u>**equencing (SPOTS), a novel multimodal approach that enables simultaneous recording of whole transcriptomes and a large panel of extracellular proteins in intact tissues (**Fig. 1a; Supplementary Table 1**). By leveraging the poly-A capture technology of the Visium slide^10^ (**Supplementary Fig. 1a**), SPOTS can simultaneously measure extracellular protein levels via polyadenylated antibody-derived tag-conjugated (ADT-conjugated) antibodies, as in CITE-seq^11^ (**Supplementary Fig. 1b**) and mRNA expression while preserving tissue architecture (**Supplementary Fig. 1c,d**). We demonstrated that by supplementing Visium’s transcriptome profiling with >30 surface markers, SPOTS yields a more granular landscape of tissues, including cell types, biological processes, and phenotypes in a highly reproducible manner.

**Figure 1:**
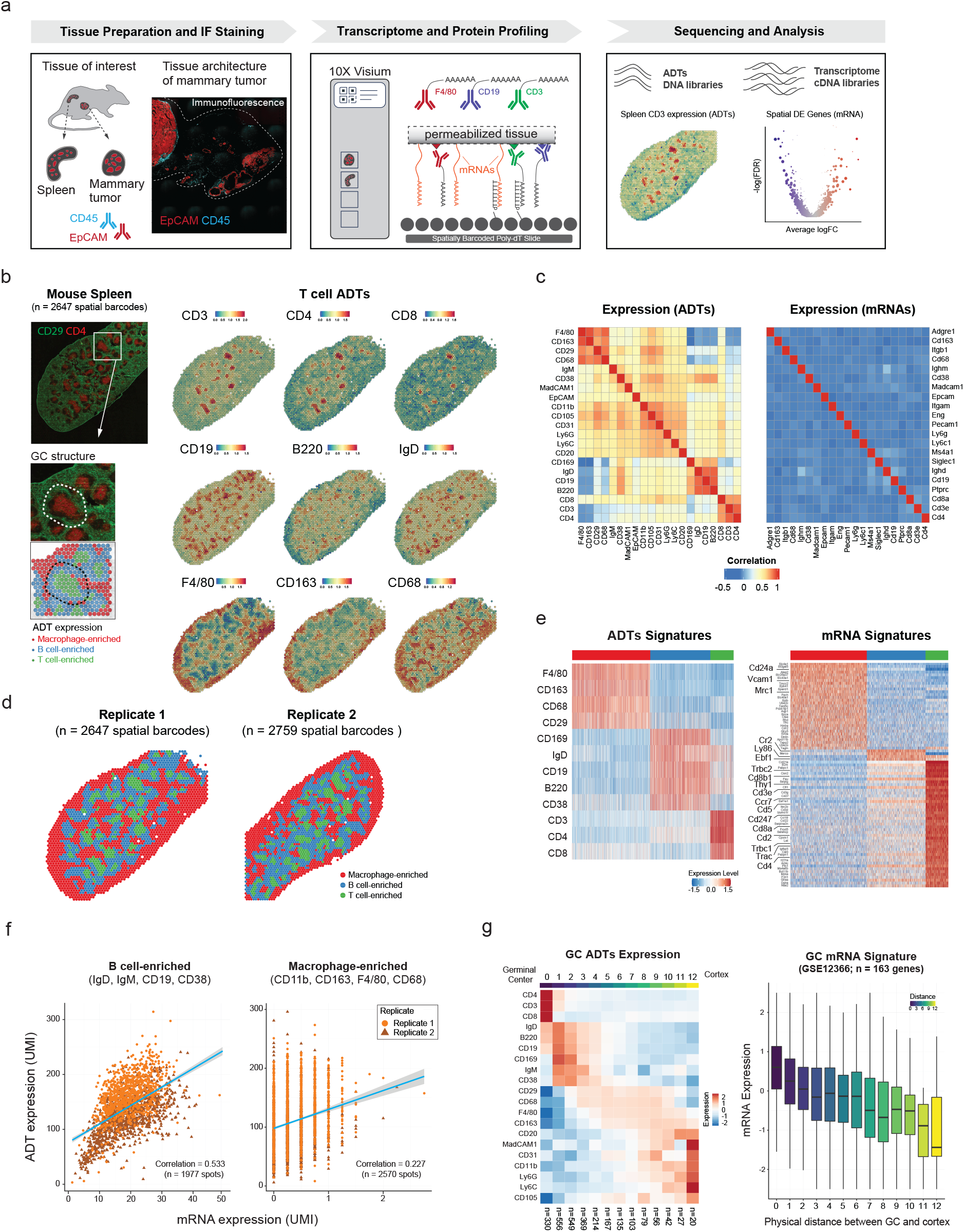
<u>S</u>patial <u>P</u>r<u>O</u>tein and <u>T</u>ranscriptome <u>S</u>equencing (SPOTS). **(a)** Overview of SPOTS workflow outline. Fresh tissue samples were collected and embedded in OCT. Tissue sections (10μm) were processed following SPOTS protocol (including staining with fluorescent and ADT antibodies, permeabilization, mRNA capture, reverse transcription, and second-strand synthesis; **Methods**) and sequenced for downstream analysis. **(b)** Normalized ADT levels of key surface markers for mouse spleen. **Left panel:** immunofluorescence (IF) staining for CD29 (green) and CD4 (red) with inset showing the germinal center (GC) architecture and its underlying spatial barcodes. **Right panel:** normalized ADT levels of marker genes for T cells (CD3, CD4, CD8), B cells (CD19, B220, IgD), and Macrophages (F4-80, CD163, CD68). **(c)** Correlation between the 21 ADTs and their corresponding mRNA expression levels (**Supplementary Table 1**) across two biological replicates. **(d)** Spatial barcode clustering and major cell type enrichment (Macrophage, B cell, and T cell) based on ADTs of two biological replicates (**Supplementary Table 2**). **(e)** ADT and mRNA signatures for each cluster of spatial barcodes. **Left panel:** heatmap showing expression levels (Z-score) of differentially expressed ADTs for each cluster. **Right panel:** heatmap showing expression levels (Z-score) of differentially expressed mRNAs in 3000 downsampled spatial barcodes. Key marker genes were highlighted (**Supplementary Table 3**). **(f)** Correlation between mRNA and ADT levels (UMI) at single spatial barcode level for B cell- and macrophage-enriched clusters in two biological replicates. **(g)** Spatial gene expression patterns of ADTs and mRNA in the spleen. **Left panel:** spatial expression pattern of the ADTs (Z-score) from GC to spleen cortex. The top color bar represents the physical distance as in the right panel. The number of spatial barcodes in each bin is labeled at the bottom. **Right panel:** boxplot showing mRNA expression levels of germinal center (GC) specific genes (GSE12366; n = 163 genes) as a function of the distance from the center of GCs. The boxes were colored by the physical distance from the center of GCs as shown in **Supplementary Fig. 3c**.

First, we optimized the assay on splenic tissues as they contain diverse immune cell populations that compartmentalized in well-defined structures of the germinal centers (GCs)^12^ (**Fig. 1b**). To achieve maximal retrieval of mRNAs and ADTs, we examined several methods of tissue fixation, antibody staining, and permeabilization conditions (**Methods**). The standard Visium workflow requires the use of methanol (MeOH) as fixative which is optimal for mRNA capture. Since MeOH fixation is not compatible with staining of most antibody clones, we optimized the assay with paraformaldehyde fixation, which is optimal for immunostaining.

We initially observed low cDNA yields and poor antibody signal (**Supplementary Fig. 1e**) and hypothesized that the tissue permeabilization enzyme used as part of the Visium Gene Expression kit (10x Genomics) workflow was not sufficient to release PFA-fixed mRNA and ADT molecules for capture on the slide. Therefore, we increased the permeabilization strength by using the tissue removal enzyme from the Visium Tissue Optimization kit (10x Genomics) in combination with 1% sodium dodecyl sulfate (SDS). These technical modifications increased the yields of both mRNA and ADT (**Supplementary Fig. 1e**).

Finally, to evaluate ADT binding specificity, we immunostained spleens with dual-labeled ADT- and fluorophore-conjugated antibodies for CD4 and CD29 to highlight the CD4+ follicular T helper cells (Tfh) and CD29+ stromal cells^12^. We detected significant ADT binding in areas without tissue (**Supplementary Fig. 1f**), which we hypothesized is due to non-specific binding via the ADT’s poly-A tail to the slide surface. To circumvent this potential non-specific binding, we added poly-dT oligos into the staining buffer to inhibit ADTs binding to surface spots, which ultimately eliminated binding on non-tissue regions and increased the signal-to-noise ratio (**Supplementary Fig. 1f**).

We next assessed SPOTS’ ability to capture cellular heterogeneity in the murine spleen using the ADT-conjugated antibodies. We designed a 21-plex panel that allows identification of cellular constituents including B cells (CD19, CD20, CD220, IgD, IgM, CD38), T cells (CD3, CD8, CD4), and macrophages (CD169, F4/80, CD163, CD68, CD11b) (**Supplementary Table 1**). Importantly, the ADT levels recapitulated the expected spleen structure^13^ as revealed by CD4 and CD29 fluorescent signals, showing CD4+ T cell-enriched naive GCs and CD29+ stroma-enriched cortex (**Fig. 1b**). Furthermore, we observed a high reproducibility (Pearson’s *r* = 0.998, p-value < 2.2e-16) in both the ADT-conjugated antibodies and mRNA expression across two independent biological replicates (**Supplementary Fig. 3a**).

Remarkably, the protein co-expression patterns showed a stronger correlation structure compared to the corresponding mRNAs (**Fig. 1c**). For instance, ADT markers of T cells (CD3, CD4, CD8) were positively correlated with each other while negatively correlated to macrophage markers (F4/80, CD163, CD68). However, such covariations between key cell-type marker genes were lacking in mRNA (**Fig. 1c**), this is in agreement with the original CITE-seq study that the ADTs are less prone to ‘dropout’ events compared to mRNAs^11^. Accordingly, we leveraged protein expression to cluster the spatial barcodes into phenotypically distinct groups that were enriched in varying degrees of macrophages, B cells, and T cells (**Fig. 1d**). In general, cell-type specific mRNAs, such as T-cell receptor-related genes (Cd3e, Cd4, Cd8a, Trac, Trbc1/2, Ccr7, Lat), were efficiently captured, as well as the corresponding surface protein counterparts (CD3, CD4, CD8; **Fig. 1e**). Surface protein and mRNA abundance also showed a positive correlation (Pearson’s *r* = 0.53, p-value < 2.2e-16) in B cell-enriched regions in which key marker genes (IgD, IgM, CD19, CD38) were well expressed in both protein and mRNA (140±0.97 ADT UMIs, 19±0.15 mRNA UMIs; mean±s.e.m) (**Fig. 1f** and **Supplementary Fig. 3b**). However, such agreement declined in macrophages-enriched regions (Pearson’s *r* = 0.23, p-value < 2.2e-16) in which key marker genes (CD11b, CD163, F4/80, CD68) were more lowly expressed in mRNA (109±0.91 ADT UMIs, 0.36±0.01 mRNA UMIs; mean±s.e.m), as previously reported^11,15^.

Next, we incorporated spatial information into our analysis to identify spatial protein and gene expression patterns within the mouse spleen. Notably, GC areas were enriched with T cells (CD3, CD4, CD8) and surrounded by B cells (IgD, IgM, B220, CD19, CD20) and CD169+ antigenpresenting macrophages, as expected in spleens of unstimulated mice^13^ (**Fig. 1g**). Further analysis showed that GC-specific genes (n = 163 genes, GSE12366) exhibited an inverse relationship with physical distance from the center of GCs (**Fig. 1g** and **Supplementary Fig. 3c**). These results highlight that by combining the protein and mRNA modalities, SPOTS provided more detailed spatial molecular profiles of cellular phenotypes than either modality alone.

Unlike spleen tissues where immune cells are highly abundant and compartmentalized, the tumor microenvironment (TME) is highly variable and complex, leading to difficulties in characterizing less abundant cell types (e.g. T cells; **Supplementary Fig. 4a**) with current ST technologies^16^. Although several single-cell RNA-seq analyses of breast carcinomas have provided a granular characterization of the tumor-infiltrating leukocytes^2,17,18^, the precise location of these signatures and their physical relation to each other are incompletely resolved. Therefore, we applied SPOTS to murine breast tumors, utilizing the MMTV-PyMT transgenic mouse model^19^ (**Fig. 2a**). To adapt the method for highly fibrotic tissue such as solid tumors, we modified the above described SPOTS tissue permeabilization protocol, thereby increasing the generalizability of SPOTS (**Fig. 2a** and **Supplementary Fig. 4b**; **Methods**). To capture the full spectrum of the immune landscape in tumors, we designed a 32-plex ADT-conjugated antibody panel to identify major cell-lineages and activation states, including tumor epithelial cells (EpCAM, KIT), fibroblasts (PDGFRA, PDPN), myeloid cells (CD11b, CD11c, F4/80, MHC-II, Ly6G, Ly6C), lymphoid cells (NK1.1, B220, CD4, CD8), and endothelial cells (CD31, TIE-2) (**Supplementary Table 1**).

**Figure 2:**
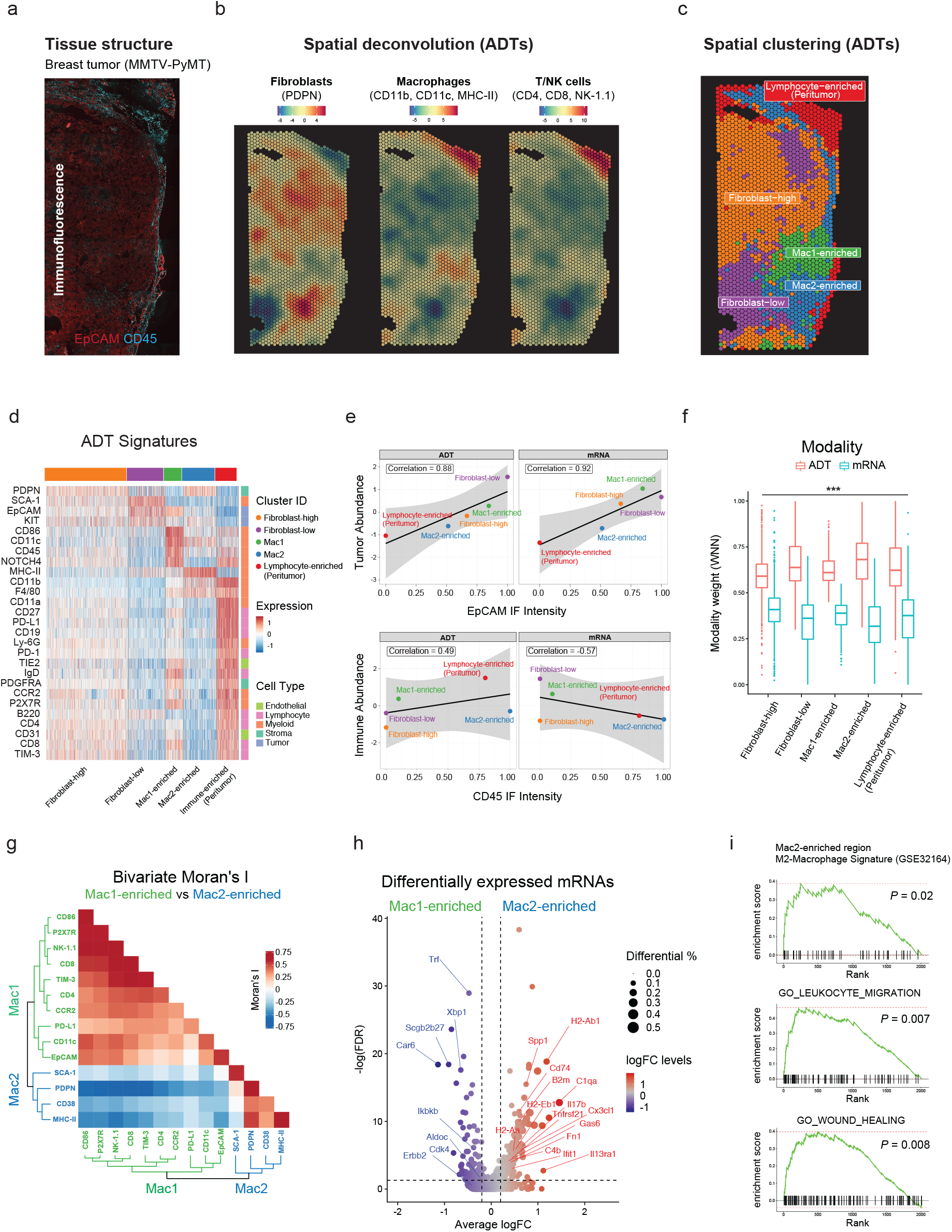
SPOTS reveals two spatially distinct macrophages in breast cancer TME. **(a)** Immunofluorescence (IF) staining of EpCAM (red) and CD45 (cyan) in breast cancer tissue. **(b)** Cell-type deconvolution based on ADTs (Z-score) of each spatial barcode overlaid onto the breast cancer tissue. Deconvolution values are provided in **Supplementary Table 4**. **(c)** Spatial barcode clustering results with annotation of major cell type enrichment based on ADTs (**Supplementary Table 4**). **(d)** Heatmap showing ADT signatures (Z-score) for each cluster (**Supplementary Table 5**). **(e)** Correlation between deconvoluted cell-type abundance (Z-score) and IF intensities across mRNA and ADT modalities at the cluster level. Pearson’s correlation coefficients were shown in the boxed labels. **(f)** Boxplot showing relative contributions (weights) to spatial barcode clustering of mRNA and ADT modalities. Paired t-test, *p < 0.05, **p < 0.01, ***p < 0.001, otherwise not significant (n.s). **(g)** Heatmap showing the spatial correlation (bivariate Moran’s I) of ADTs in Mac1- and Mac2-enriched regions with dendrograms showing hierarchical clustering of ADTs. **(h)** Volcano plot showing log fold changes (logFC) of the top 5000 most variable genes between Mac1- and Mac2-enriched spatial barcodes and their significance (y-axis; −log10 scale). Genes are dotted and colored by logFC levels (color scale). The size of the dot represents the difference in the fraction of detection between the two groups. Macrophage-related genes are annotated. P-values were determined by the Wilcoxon Rank Sum test. Vertical dotted lines represent ±0.2 logFC. Horizontal dotted lines represent false discovery rate (FDR) of 0.05 (-log10 scale). Raw and FDR corrected P-values and logFC values are listed in **Supplementary Table 5**. **(i)** Gene Set Enrichment Analysis (GSEA) of Mac2-enriched spatial barcodes based on mRNA expression.

In order to characterize the complex interactions and the spatial gene expression heterogeneity within the TME, we first performed ADT-based cell-type deconvolution^20^ and clustering^21^ analyses (**Fig. 2b,c; Methods**). This revealed several distinct tumor and peritumor regions enriched with varying degrees of immune and stromal cells (**Fig. 2d** and **Supplementary Fig. 5a**). As observed in the splenic tissue (**Supplementary Fig. 2b**), the correlation between the ADT and its corresponding mRNA was high when the gene was well expressed (**Supplementary Fig. 5b**). By further comparing the ADT- and mRNA-based cell-type deconvolutions, we observed a positive correlation between tumor cell abundance and the corresponding EpCAM immunofluorescence (IF) signals in both ADT- (Pearson’s *r* = 0.88) and mRNA-based (Pearson’s *r* = 0.92) deconvolutions (**Fig. 2e**). However, the correlation between immune cell abundance and the corresponding CD45 IF signals was poor in mRNA-based deconvolution (Pearson’s *r* = −0.57) compared to ADT-based deconvolution (Pearson’s *r* = 0.49), likely due to low abundance of immune-related mRNAs (**Supplementary Fig. 5c**). Indeed, by using a weighted nearest neighbor (WNN) based multimodal integration method^21^, we found that the ADT modality had a greater impact on clustering assignment compared to mRNA (**Fig. 2f**; 0.63 vs 0.37 WNN weights, 95%CI of mean differences 0.25-0.27, paired t-test p-value < 2.2e-16). These results further reinforced that by integrating both the ADT and mRNA modalities, SPOTS provided a more precise molecular characterization of immune cell populations in disorganized tissues, such as solid tumors.

We next examined two regions that were enriched with macrophage/myeloid markers (Mac1- and Mac2-enriched; **Fig. 2c,d**). Phenotypically, Mac1-enriched regions exhibited immunostimulatory ADTs of myeloid cells (CD86, CD11c) while Mac2-enriched regions displayed immunosuppressive ADTs (PD-L1; **Fig. 2g**). Structurally, the Mac2-enriched areas were in close proximity to a lymphocyte-enriched region (**Fig. 2c**) that was enriched with T cell exhaustion markers (PD-1, TIM-3), in-line with the well documented immunosuppressive function of tumor-associated macrophages^19^. These results were further validated by differential mRNA expressions between Mac1- and Mac2-enriched spatial barcodes (Spp1, C1qa; **Fig. 2h** and **Supplementary Fig. 5d**), and gene set enrichment analyses for M2-macrophages and wound healing signatures (**Fig. 2i**). These data suggest that M2-macrophages are forming an immunosuppressive barrier at tumor borders (**Fig. 2c**), leading to immune exhaustion and evasion, as previously shown in breast cancer^18^.

Here, we present SPOTS, a method that allows simultaneous measurement of a large panel of protein markers and whole transcriptomes in intact tissues. We envision that SPOTS–a technique that makes use of the broadly available Visium technology and polyadenylated DNA-barcoded antibodies–may be readily integrated into emerging higher resolution ST technologies^4,5^, potentially further improving the molecular characterization of cell types and states in tissues. Future developments in SPOTS to capture somatic mutations in tumor cells^22^ or TCR sequences of tumor-infiltrating T cells^23^, would also enhance our understanding of cell-cell interactions in the immune microenvironment.

Collectively, our data across normal spleen and breast tumors demonstrated that SPOTS enables both in-depth gene expression analysis and highly multiplexed immunophenotyping, and is highly reproducible, simple to execute, and compatible with commercially available platforms. Broad adoption of SPOTS by the genomics community will pave the way towards a spatially-resolved multimodal map of tissue organization and function in health and disease.

## Methods

### Mouse work

Animal procedures were approved by the Institutional Animal Care and Use Committee (IACUC) of the Research Animal Resource Center (RARC) at Weill Cornell Medicine (Protocol: 2016-0058). All mouse strains involved in this work including MMTV-PyMT (Stock No: 022974) and wild-type C57BL/6J (Stock No: 000664) were purchased from The Jackson Laboratory (USA). Females of the MMTV-PyMT model developed spontaneous tumors after ~110 days post-birth. To harvest fresh tumors, mice were euthanized and mammary gland tumors were collected, washed three times in PBS-1% BSA, and embedded in OCT for storage at −80C.

### List of antibodies used for SPOTS

The polyadenylated Biolegend TotalSeq™ A anti-mouse antibodies and clones used were: CD4 (Clone RM4-5); CD8a (Clone 53-6.7); CD366 (Tim-3) (Clone RMT3-23); CD279 (PD-1) (Clone RMP1-30); CD117 (c-kit) (Clone 2B8); Ly-6C (Clone HK1.4); CD11b (Clone M1/70); Ly-6G (Clone 1A8); CD19 (Clone 6D5); CD45 (Clone 30-F11); CD25 (Clone PC61); CD45R/B220 (Clone RA3-6B2); CD11c (Clone N418); F4/80 (Clone BM8); I-A/I-E (Clone M5/114.15.2); NK-1.1 (Clone PK136); Ly-6A/E (Sca-1) (Clone D7); CD3 (Clone 17A2); CD274 (B7-H1, PD-L1) (Clone MIH6); CD27 (Clone LG.3A10); CD20 (Clone SA275A11); CD86 (Clone GL-1); MadCAM1 (Clone MECA-367); CD163 (Clone S15049I); CD192 (CCR2; Clone SA203G11); CD169 (Siglec-1) (Clone 3D6.112); CD326 (Ep-CAM) (Clone G8.8); IgM (Clone RMM-1); CD38 (Clone 90); CD68 (Clone FA-11); CD29 (Clone HMβ1-1); IgD (Clone 11-26c.2a); CD140a (Clone APA5); CD11a (Clone M17/4); CD105 (Clone MJ7/18); P2X7R (Clone 1F11); CD1d (CD1.1, Ly-38) (Clone 1B1); Notch 4 (Clone HMN4-14); CD31 (Clone 390); CD202b (Tie-2, CD202) (Clone TEK4); Podoplanin (Clone 8.1.1). See **Supplementary Table 1** for a complete list of antibodies, clones, and barcodes used for SPOTS.

### Fluorophore conjugation of ADT-conjugated antibodies

Biolegend TotalSeq™ A anti-mouse antibodies CD4 (Clone RM4-5) and CD29 (Clone HMβ1-1) were conjugated with Alexa 555 or Alexa 647 with the Antibody Labeling Kit (ThermoFisherScientific) according to manufacturer instructions.

### Pre-SPOTS quality control and optimization

In advance of SPOTS, sections were obtained for RNA quality measure with the RNeasy Plus kit. Tissues with an RNA integrity number (RIN) of 7 or greater were used for future experiments. The optimal ADT concentrations were determined by titrations of fluorescently-tagged matched clones using immunofluorescence (IF). Appropriate permeabilization conditions for both the pre-staining permeabilization and the full tissue permeabilization were determined using the 10x Genomics Tissue Optimization kit (**Supplementary Table 1**).

### Solutions for SPOTS

The staining and blocking solutions were adapted as previously described^9^. The following solutions were made immediately in advance of the experiment and were kept on ice. **2x blocking solution** (75μl per sample): 6x SSC, 0.2% Tween 20, 4% BSA, 0.2μg/μl sheared salmon sperm, 2.0 U/μl RNAse inhibitor. **Pre-staining permeabilization solution**: 0.25% saponin (5x), (Thermo Fisher J63209.AK), 0.4U/μl RNAse inhibitor, remaining volume is compensated with PBS. **Antibody staining solution** (50μl per sample): 1x blocking buffer (dilution of 2x blocking buffer above), 20μM blocking oligos (dT25), fluorescent and TotalSeq™ A (Biolegend, USA) antibodies as titrated (**Supplementary Table 1**); 1x blocking buffer (75μl per sample): 1x blocking buffer, 5mM ribonucleoside vanadyl complex (heated to 65C shortly before use), Fc block (BioLegend, USA; 5μl per sample); wash buffer (600μl per sample): 1x blocking buffer. Any remaining volume was compensated with water unless noted otherwise.

### SPOTS

10μm thick, OCT-embedded sections were deposited on the 10x Genomics Visium slides; slides were stored at −80C for up to 1 week in advance of the experiment. Slides were moved on dry ice to the thermocycler and then incubated for 1 minute at 37C on the Thermocycler Adaptor. The slides were then immersed in 1% PFA in PBS for 10 minutes at room temperature to fix the tissue. The slides were washed by dunking five times into a falcon tube with 3x SSC and then secured in the cassette. Each sample was blocked in 75μl of 1x blocking buffer for 15 minutes at 4C. Post-blocking, 75μl of the pre-staining permeabilization solution was added to each sample at 4C for 10 minutes. The solution was removed, and 50μl of antibody staining mix was added to each sample. The slide was then incubated and protected from light for 90 minutes at 4C with the antibodies. The samples were then washed with 100μl of wash buffer four times for one minute each. A final pre-imaging wash was conducted by dunking the slide twenty times in a falcon tube with 3x SSC. The slide was imaged at 10X magnification to detect APC (CD45) and PE (EpCAM) fluorophores using the Axio Observer 7 Inverted Microscope (Zeiss Microscopy).

After imaging, the tissue was fully permeabilized for 9 minutes for spleen tissue and 27 minutes for breast cancer tissue using the Tissue Optimization Tissue Removal Enzyme (per sample: 3.5μL Tissue Removal Enzyme (10x Genomics, 3000387), 59.5μl 1x SSC, 7μl 10% SDS) **(Supplementary Fig. 4b)**. Each well was then washed three times with 150μl of 0.1x SSC. The remainder of the experiment was conducted using the 10x Genomics Gene Expression protocol, starting with step 2.2 Reverse Transcription, including Visium Spatial Gene Expression Library Construction. The only deviations from the original protocol were the addition of additive primer to amplify ADT during Second Strand Synthesis (Step 3.0) and cDNA Amplification (Step 4.2) (8.8μl of 100uM and 4μl of 0.2μM additive primer were added to each reaction, respectively) (**Supplementary Table 1**) and retention of the supernatant in step 4.3 cDNA Cleanup - SPRIselect. Two sequential 1.9x SPRIselect cleanups (130μl and 95μl of SPRI for the first and second cleanups, respectively) were performed on said supernatant and the final product was eluted in 45μl of water. The ADT product from the supernatant was amplified and indexed using the SI-PCR and TruSeq Small RNA RPIx primers before Illumina sequencing (**Supplementary Table 1**). The master mix and thermocycler conditions for this reaction were as follows: 45μl ADT, 50μl of Amp Mix (2000047), 2.5μl of 20μM TruSeq Small RNA RPI index primer, 2.5μl of 20μM SI-PCR primer; 95C for 3 minutes > (95C for 20 seconds, 60C for 30 seconds, 72C for 20 seconds, 10x cycles) > 75C for 5 minutes > 4C hold. Approximately 50,000 and 8,500 reads were designated per spatial barcode for gene expression and ADT sequencing, respectively. Transcript (GEX) and antibody derived tag (ADT) libraries were sequenced on the Illumina Nextseq500 (spleen) or Novaseq6000 (mouse breast cancer).

### Spleen data analysis

The sequencing data and histology images were processed using SpaceRanger (v1.3.0 and pre-release v.2.0, 10x Genomics) with manual tissue alignment using Loupe Browser (v5.0, 10x Genomics) to obtain raw UMI count spot matrices for ADT and GEX. We analyzed the data using Seurat 4.0^21^. Briefly, ADT data from the two spleen replicates was normalized using CLR normalization across each cell with Seurat’s *NormalizeData* function, and the transcriptomic data was normalized using log-transformed transcript per million (logTPM). We scaled and centered the normalized ADT data and then performed Principal Component Analysis (PCA) using Seurat’s *RunPCA*. We selected the first 10 PCs to build a shared nearest neighbor graph using Seurat’s *FindNeighbors* function and clustered the spatial barcodes using the *FindClusters* function with *resolution=0.2*. Upon clustering, we noticed a cluster of 15 spatial barcodes out of the 5421 total barcodes, that are sporadically spread and have a significantly lower number of ADTs compared to the rest (18 vs. 1248 UMIs). We reasoned that these spatial barcodes may represent technical failures and we removed them from downstream analysis and retained 5406 spatial barcodes (2647 from Replicate 1 and 2759 from Replicate 2). There were 10 spatial barcodes near the edge of the slides that were excluded for visualization purposes. In total, we retained four clusters of spatial barcodes that were enriched in Macrophage, B, T, and epithelial cells, and we excluded the epithelial cell-enriched cluster for visualization purposes. The clustering result and annotation are listed in **Supplementary Table 2**. To visualize the sparse mRNA data, we randomly sampled 30 barcodes for each cluster and created a downsample of 3000 spatial barcodes.

### Breast cancer data analysis

Tumor tissues from the MMTV-PyMT mouse model were first aligned onto the Visium slides with Loupe Browser 5.0 (10x Genomics) using IF images as input. The resulting alignment JSON files were used as input for SpaceRanger software (v.1.3.0 10X Genomics), and the transcriptomic reads were mapped to a custom genome (mm10 with PyMT transgene). The ADT reads were mapped to the antibody barcode sequences (**Supplementary Table 1**) with the 4992 whitelist tissue spatial barcodes provided by 10X Genomics as input using the Python package CITE-seq-Count v1.4.3^24^. The count matrices of the transcriptome and ADT data were then normalized using log-transformed transcript per million (logTPM).

### Data visualization

To better visualize the SPOTS data, we utilized local *Gi\** statistics^25^, which is similar to local Moran’s I and local Geary’s C statistics^26^, to create smooth expression profiles for visualization and diagnostics. The local *Gi\** statistics for each feature/gene was defined as 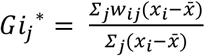 where *x_j_* is the expression value of a feature in cell *j* and *w_ij_* is the spatial weight between cell *i* and *j*; the 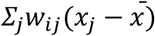 term is interpreted as weighted neighborhood or spatial lag. By definition local *Gi\** statistics is a Z-score and its *P*-value can be calculated using permutation-based methods^26^ or mathematical derivations^27^. Because Visium tissue barcodes are arranged on hexagonal lattices, we defined the weight matrix using hexagonal nearest neighbor distance (HNN)^28^, which reflects its true spatial orientation. We assigned the weights using HNN with a gaussian kernel of width 1 unit distance, and only neighbors within a 3 hexagonal distance were considered. We measured the physical distance from the center of GCs (defined by T cell-enriched regions) using HNN. The physical distance from the center of GCs for all the spatial barcodes are listed in **Supplementary Table 2**.

### Cell type deconvolution

We performed deconvolution of the breast cancer SPOTS data using SPOTlight v0.1.7 R package^20^ with default parameters, except we set *min_cont=0* which did not remove any cells, and a recent unbiased scRNA-seq data of wild type MMTV-PyMT mouse model^17^ as reference. Briefly, the scRNA-seq data were downloaded from GSE158677 and analyzed using Seurat 4.0^21^. Upon clustering, we identified six major cell types including Tumor, CAF, Endothelial, Myeloid, B and T/NK cells. Differentially expressed genes were calculated by Seurat’s *FindAllMarkers* function with parameters *min.pct=0.5* and *logfc.threshold=1* and used for deconvolution with SPOTlight’s *spotlight_deconvolution* function. For the ADT deconvolution, we first scaled the cell type ADTs (Tumor: EpCAM and KIT; CAF: PDPN; Macrophage: CD11c, CD86 and F4/80; B cell: CD19 and B220; T/NK: CD4, CD8 and NK-1.1) using cosine normalization^29^ and calculated the mean expression of their ADTs for each cell type. The deconvoluted percentages were calculated as the fraction of the cell type ADT expressions over the total sum of all ADTs. The scaled deconvolution values were calculated using local *Gi\** statistics as described above. The IF staining intensity was quantified for each spatial barcode and normalized to the 0-1 range, outlier values below 1st or above 99th percentiles were removed. The deconvolution values for each barcode are listed in **Supplementary Table 3**.

### Integration

In order to integrate the transcriptome and ADT data and assess the contributions of each modality, the WNN weights were calculated using Seurat’s *FindMultiModalNeighbors* function using the first 30 PCs of transcriptome and first 10 PCs of ADTs. The WNN weights of mRNA and ADT modalities are listed in **Supplementary Table 3**.

### Spatial cross-correlation analysis

The spatial cross-correlation of each gene can be defined by either univariate or bivariate Moran’s I. In the univariate case, Moran’s I of any feature *X* was calculated as 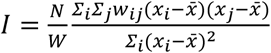, where *w_ij_*, is a spatial weight matrix with *w_ii_* = 0 and *W* = *∑w_ij_*, and *N* is the number of total spatial barcodes. The *P*-value of univariate Moran’s I can be obtained by using mathematical derivations^30^. In the bivariate case^31^, Moran’s I between feature *X* and *Y* was calculated as 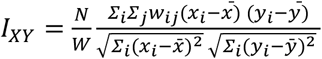, and the *P*-value can be obtained by using mathematical derivations^32^.

### Spatial clustering

To identify spatially distinct cell populations, we modified the current clustering scheme, which is solely based on expression or ADT data by incorporating the spatial information of each spatial barcode. We adopted a method that has been developed in the field of spatial statistics^31^, where multivariate spatial correlation (MSC) was defined as 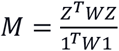 (*Z* is the z-score of the data matrix). This *M* matrix captures the spatial relationship between each gene across tissue spatial barcodes and is used for subsequent principle component analysis. This approach entails that the spatial correlation matrix (*M*) must be positive semidefinite, like the covariance matrix used in Principal Component Analysis (PCA), where the eigenvalues must be nonnegative. However, the *M* matrix is not positive semidefinite and can have negative eigenvalues as noted in the original publication. To resolve this, another spatial correlation^33^ matrix *L* with 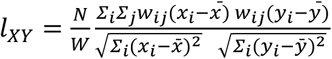, such that 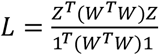, is positive semidefinite. By using this *L* matrix, we performed the singular value decomposition (SVD) in the same manner as PCA, and alternative sparse solutions can be obtained accordingly^34^. We termed this approach the Spatial Component Analysis (SCA) to imply its connection to canonical PCA. For clustering of the ADT data, we used the HNN distance weights as described in the **Data visualization** and the first 10 Spatial Components (SCs) as input to build the shared nearest neighbor graph and then clustered the spatial barcodes using the *FindClusters* function with *resolution = 0.3*. The clustering result and annotation are reported in **Supplementary Table 3**.

### Differential expression analysis

Differentially expressed (DE) genes for each cluster were calculated using Seurat’s *FindAllMarkers* function with default parameter or *logfc.threshold = 0.1* for ADT data. For DE genes between Mac1- and Mac2-enriched regions, the top 5000 variable genes with the highest dispersion values were selected using the *FindVariableFeatures* function. Gene Set Enrichment Analysis (GSEA) was performed using the *fgsea v1.16* R package^35^ and GO term biological process gene sets (c5.bp.v7.1.symbols) with 10000 permutations. M2 macrophage signatures genes were selected from GSE32164. The spleen germinal center signature genes were selected from GSE12366. The DE genes of spleen and breast cancer samples are listed in **Supplementary Table 4** and **Supplementary Table 5,** respectively.

## Acknowledgments

We acknowledge J. Chew and Y. Yin from 10x Genomics Headquarters for critical discussions and K. Ganapathy for helping coordinate 10x Genomics and Weill Cornell interactions. This work was supported by the BWF (Award #1014689.01), NCI R01 (1R01CA228135), CEGS (RM1 HG011014), and Emerson (NPT Charitable Grant).

## Data availability

All raw data generated in this study have been deposited to GEO. The accession code will be made available upon publication.

---

**Supplementary Figure 1:**
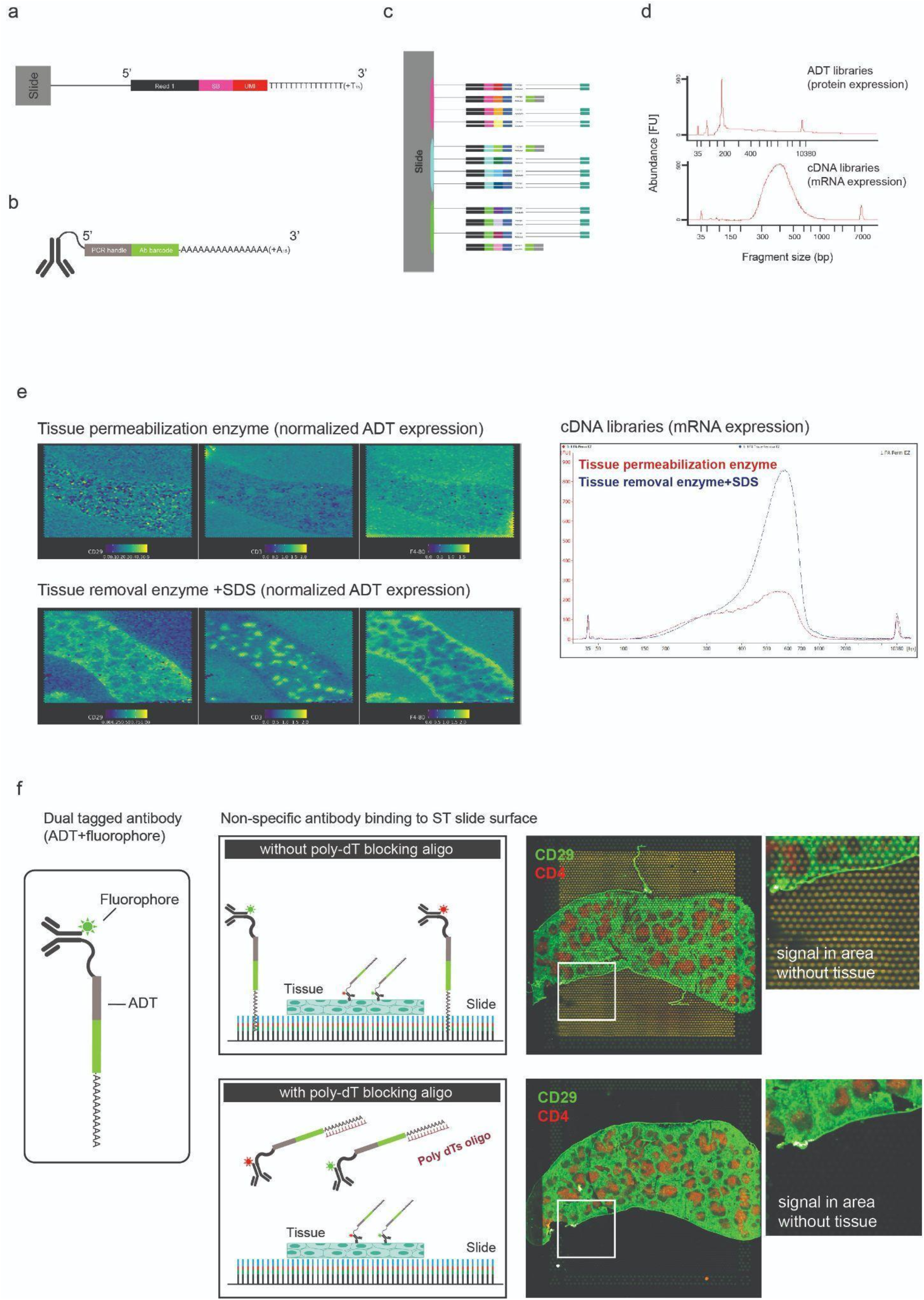
SPOTS library preparation and optimizations. **(a)** Illustration of the structure of a barcoded oligo, including Read 1, spatial barcode (SB), unique molecular identifier (UMI), and poly-T (T_30_) sequences, attached to the Visium slide. **(b)** Illustration of the structure of an ADT-conjugated antibody, including a PCR handle (Handle), an antibody barcode (AB), and poly-A (A_30_). **(c)** Schematic of SPOTS following second strand synthesis. ADT and mRNA location is recorded through spatial barcodes on the Visium slide. **(d)** After cDNA amplification, ADT and cDNA libraries can be separated and prepared for sequencing to produce indexed ADTs (top panel, BioA) and gene expression libraries (bottom panel, BioA). **(e)** Normalized ADT levels (left) and cDNA libraries (right, BioA) of indicated antibodies using tissue permeabilization vs. tissue removal enzyme and SDS. **(f)** Immunofluorescence using dual-tagged antibodies (ADT+fluorophore) of CD29 (green) and CD4 (red) in spleens in the presence or absence of poly-dT blocking (20μM). Note for reduction of non-specific bindings in non-tissue areas.

**Supplementary Figure 2:**
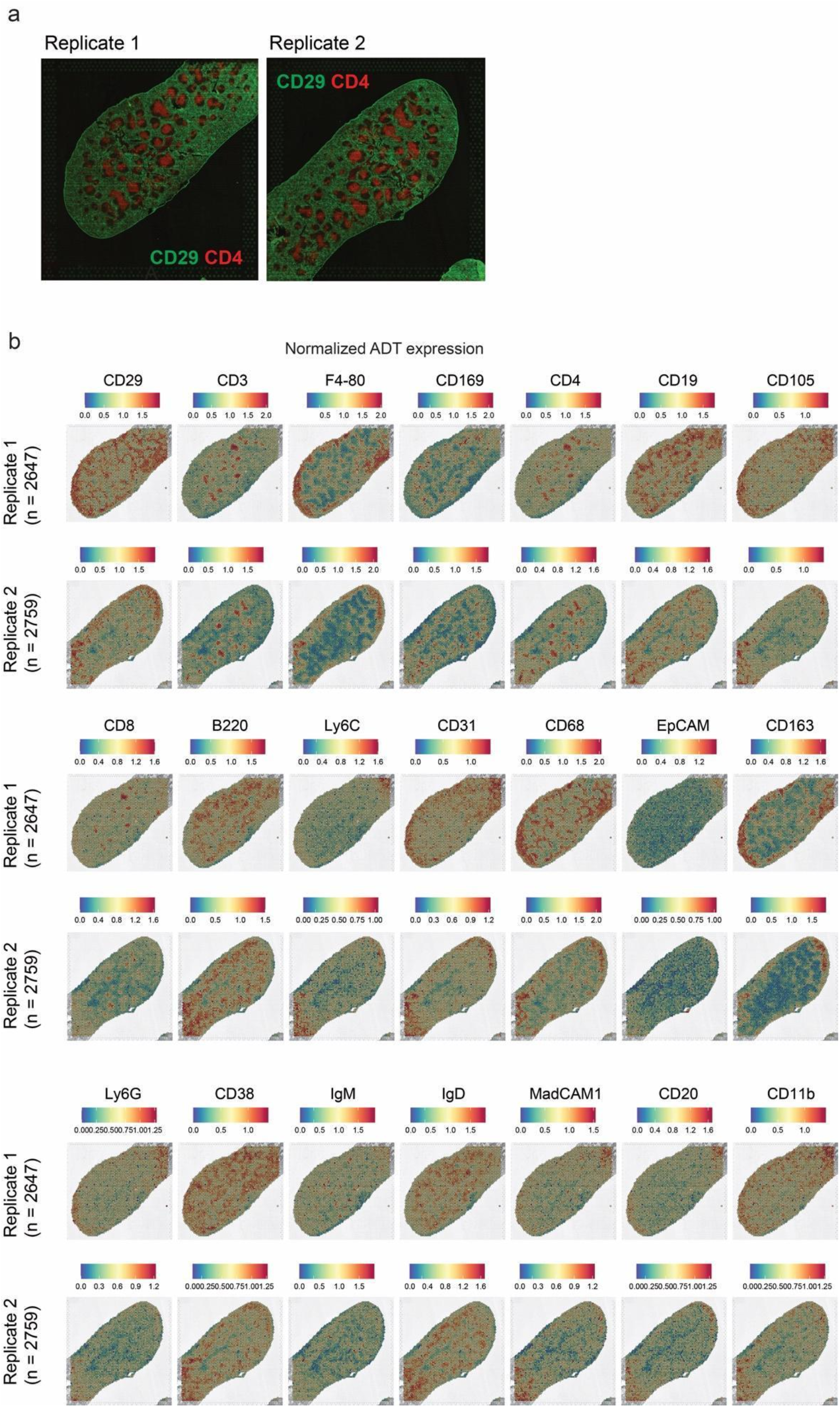
Normalized ADT levels in spleens. **(a)** Immunofluorescence staining for CD29 (green) and CD4 (red). **(b)** Normalized ADT levels of all 21 surface proteins in two spleen samples.

**Supplementary Figure 3:**
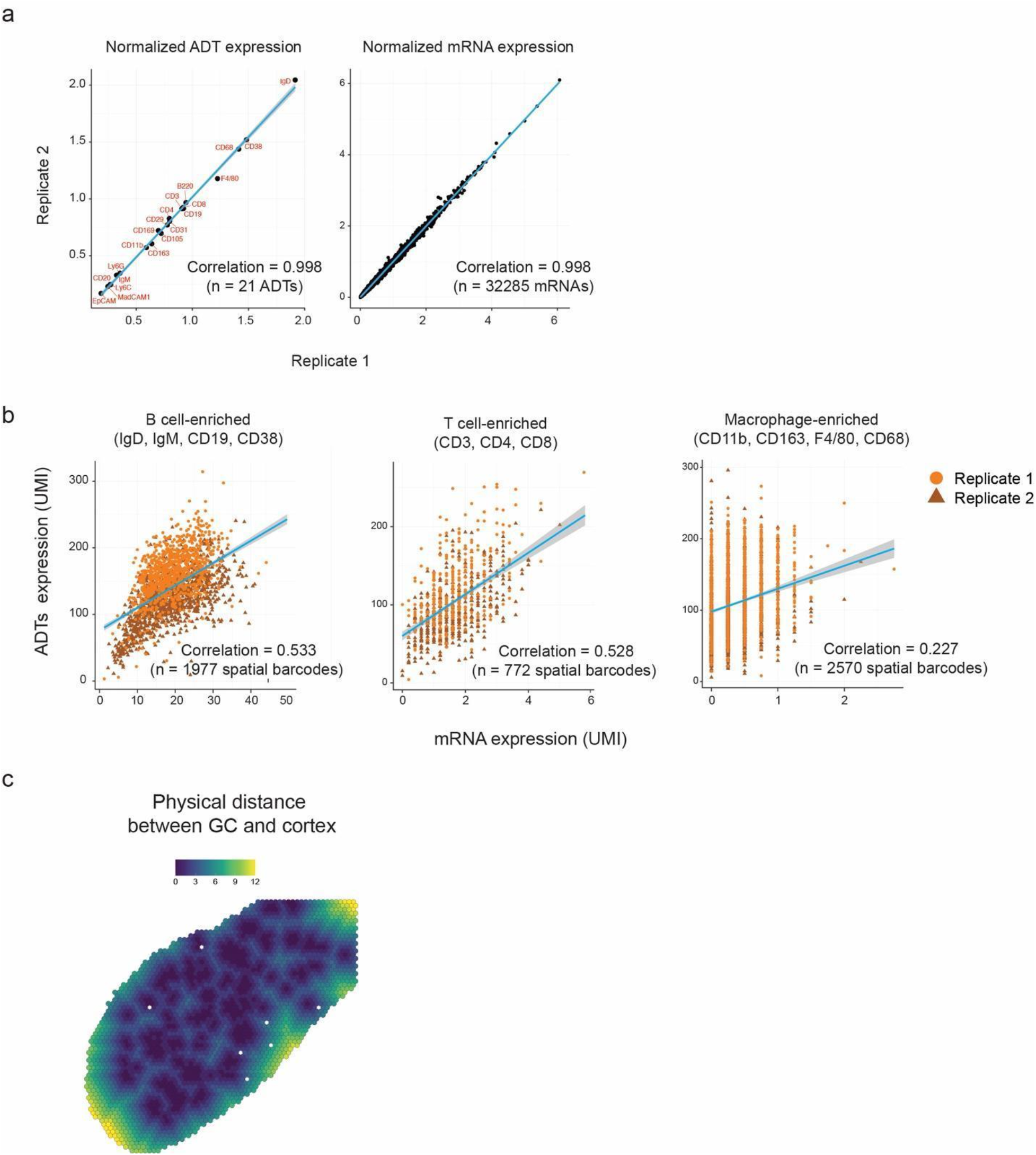
Reproducibility and quality controls of SPOTS spleen data. **(a)** Correlation of normalized ADT levels and mRNA expressions between two biological replicates of mouse spleen. **(b)** Correlation between mRNA and different ADT levels (UMI) at single spatial barcode level for B cell, T cell, and macrophage enriched clusters across two biological replicates. **(c)** Physical distance (color scale) from the center of GCs for each spatial barcode.

**Supplementary Figure 4:**
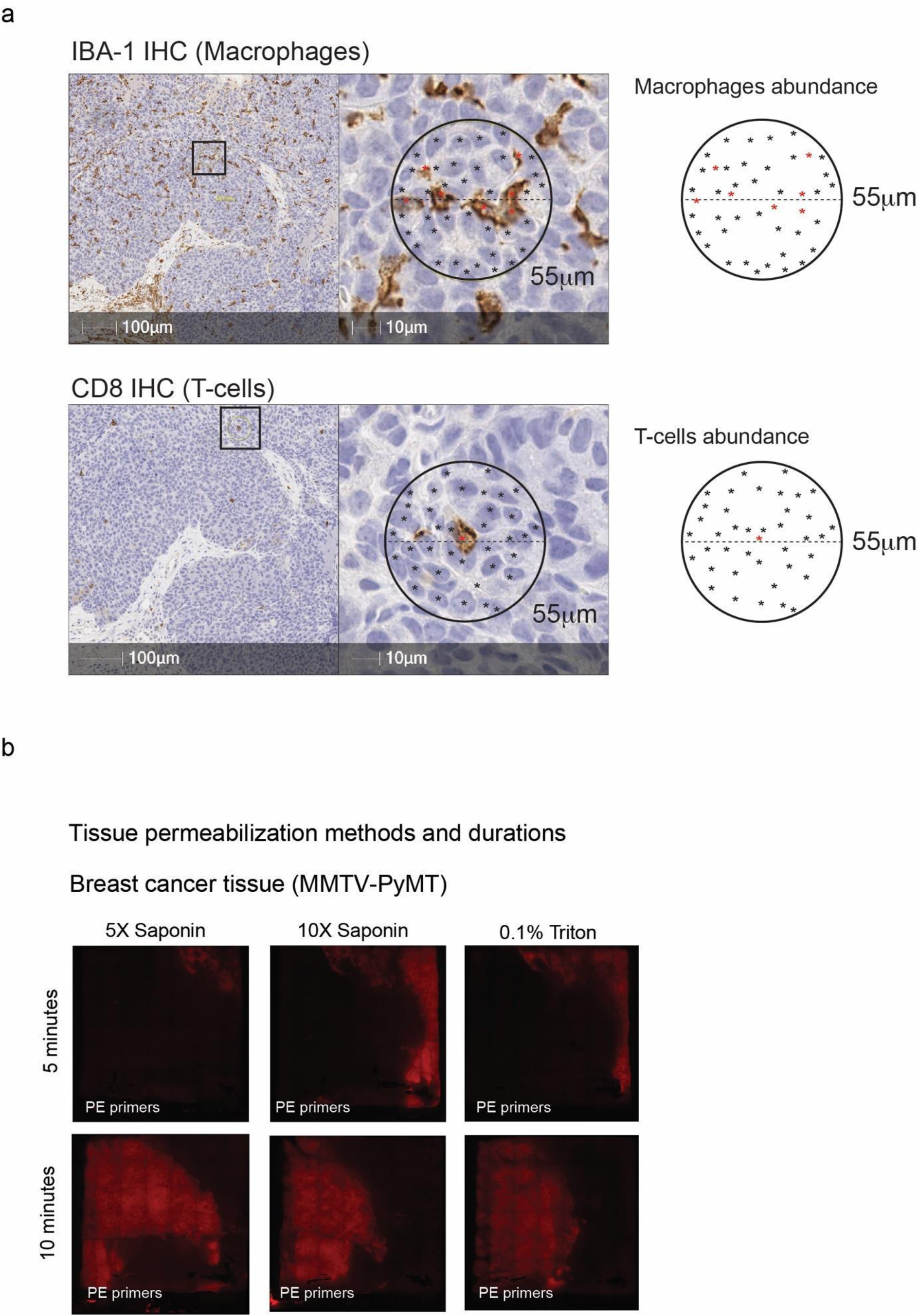
Breast cancer tissue architecture and optimizations. **(a)** Immunohistochemistry (IHC) analysis of CD8 (T cells) and IBA-1 (macrophages) in mammary tumor section from MMTV-PyMT model demonstrating the typical abundance of T cells and tumor-associated macrophages in breast cancer. **(b)** Fluorescence imaging of TRITC labeled cDNA from mouse breast cancer following 5 or 10 minutes with 5X or 10X saponin or 0.1% Triton pre-staining permeabilization conditions.

**Supplementary Figure 5:**
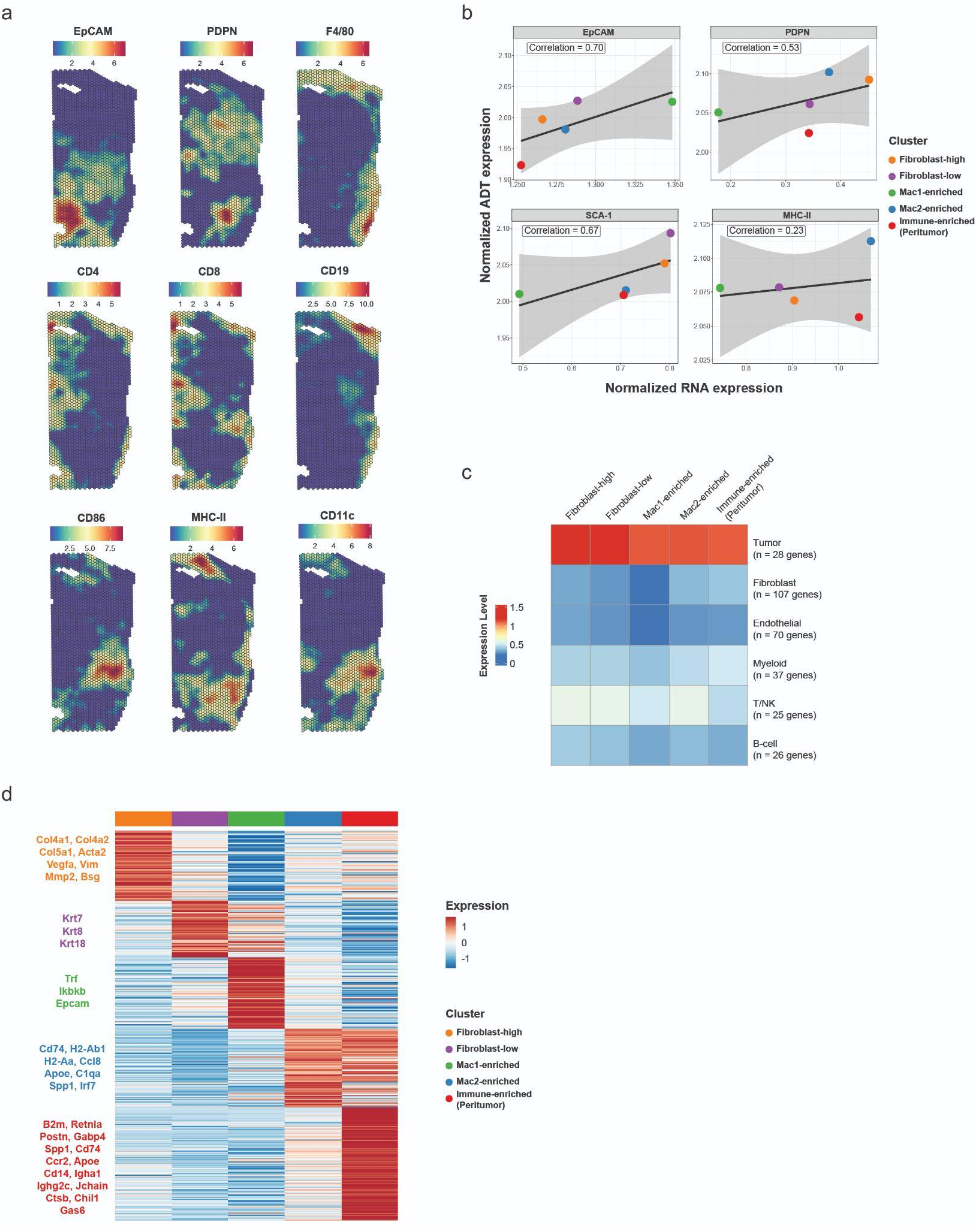
Spatial mRNA and ADT levels in breast cancer TME. **(a)** Normalized ADT levels of selected markers in MMTV-PyMT breast cancer tissue. **(b)** Correlation between normalized mRNA expression and ADT levels of the indicated surface markers (EpCAM, PDPN, SCA-1, MHC-II). Pearson’s correlation coefficients were calculated on cluster-level and are indicated in the boxed labels. **(c)** Average normalized mRNA expressions of key cell-type marker genes (GSE158677) in each spatial cluster. **(d)** Heatmap showing the average expression levels (Z-score) of differentially expressed mRNAs for each cluster. Key marker genes are colored and highlighted.

